# Membrane-bound Guanylyl Cyclase COP5/HKR1 changes ciliary beat pattern and biases cell steering during chemotaxis in *Chlamydomonas reinhardtii*

**DOI:** 10.1101/2024.10.14.618206

**Authors:** Alexis Strain, Nathan Kratzberg, Dan Vu, Emmaline Miller, Ken-ichi Wakabayashi, Adam Melvin, Naohiro Kato

## Abstract

This study investigates the control of ciliary beat patterns during ammonium chemotaxis in the model ciliate microalgae *Chlamydomonas reinhardtii*. Screening the chemotaxis response of mutant strains with ciliary defects revealed that a strain lacking CAV2, the alpha subunit of the voltage-gated Ca^2+^ channel, is deficient in ammonium chemotaxis. CAV2 regulates the switching of the ciliary beat pattern from the asymmetric to the symmetric waveform. Strains lacking COP5/HKR1 (chlamyopsin 5/histidine kinase rhodopsin 1), a membrane-bound guanylyl cyclase, are also deficient in ammonium chemotaxis. Conversely, strains defective in phototaxis perform ammonium chemotaxis normally.

Cell motility analysis revealed wild-type cells reduce the incidences of switching the ciliary beat pattern from the asymmetric to symmetric waveform when swimming up the ammonium gradient. In contrast, the COP5/HKR1 disrupted strain does not bias ciliary beat pattern switching in the gradient. This finding reveals that a membrane-bound guanylyl cyclase plays a critical role in *Chlamydomonas* chemotaxis signaling transduction, similarly to animal chemotaxis. On the other hand, ciliary beat pattern switching induces randomized directional changes, analogous to run-and-tumble chemotaxis of bacteria and archaea. This study reveals that *Chlamydomonas* signaling transduction is similar to the eukaryotic mechanism, yet the cellular locomotion follows the bacteria and archaea mechanism.

## Introduction

Motile cilia are tubular organelles protruding from the cellular membrane of selected types of eukaryotic cells. Motile cilia exhibit ciliary beat patterns, or waveforms, that can change shape, frequency, and amplitude in response to the environment. For instance, in mammalian respiratory tract cells, the ciliary beat frequency is changed to remove particles in mucus^1,2^. In sperm, another ciliated mammalian cell, the ciliary beat pattern changes to facilitate migration, or steering, toward an egg^3^. *Chlamydomonas reinhardtii* is a biciliate green alga model for eukaryotic cilia, which has previously been used to reveal the fine structure of the cilium and ciliary dynamics, including that the outer and inner dynein arms respectively regulate ciliary beat frequency and waveform^4,5^.

Another well-documented topic in *Chlamydomonas* is phototaxis (directed movement in response to light gradients) and the resulting changes in the ciliary beat pattern required to steer cells toward or away from light. However, chemotaxis (directed movement in response to chemical gradients) is poorly characterized, with an unidentified sensory system, signaling pathway, and steering by ciliary beat pattern^6-11^. Ammonium (NH_4_^+^) is a positively charged ion and one of the strongest reported chemoattractants in *Chlamydomonas*^12-14^. NH_4_^+^ as a nitrogen source is used preferentially over nitrate (NO_3_^−^) and nitrite (NO_2_^−^) and can be directly imported into the cell using ammonium transport proteins (AMT)^15,16^. Once imported, NH_4_^+^ can be assimilated into amino acids through the glutamine and glutamate synthase pathways^17^. Notably, gametic cells will lose chemotactic sensitivity to NH_4_^+^, however, will maintain NH_4_^+^ uptake^18^. A *Chlamydomonas* strain lacking AMT4, functioning as a major NH_4_^+^ importer, performs chemotaxis normally in the vegetative cells^14^. These observations suggest the existence of an ammonium-specific metabolic-independent chemotaxis receptor.

Sensor histidine kinases are the major chemoreceptors in bacteria and archaea. The sensors are part of a two-component system comprised of multiple proteins^19^. Homologous histidine kinase proteins are found in fungi and plants as well. In both bacteria and archaea, the signal is initiated by the ligand binding domain of the methyl-accepting protein, then transferred through phosphorylation of the histidine kinase protein, also known as CheA, and the response regular protein, also known as CheY^20^. The methyl-accepting protein (MCP) where methylation and demethylation occur, functions as a feedback regulator of the histidine kinase activity^21^. In archaea, the sensor histidine kinase also transduces the phototaxis signal by forming a complex with rhodopsin (sensory rhodopsin II, SRII) in the plasma membrane (Fig. S1)^22^. Eukaryotic chemotaxis sensors are typically G-protein-coupled receptors that bind to a ligand, initiating the signal and causing a conformational change, which activates a guanylyl (or adenylyl) cyclase domain, transferring the signal to the secondary messenger cyclic GMP (or AMP)^23^.

*Chlamydomonas* possess a family of protein receptors called chlamyopsins (COPs), which include the receptors for phototaxis ChR1 (COP3) and ChR2 (COP4), encoding a rhodopsin domain^24,25^. Other proteins in the family, COP5-COP10, are also histidine kinase receptors (HKRs). Typically, HKRs encode a rhodopsin domain, histidine kinase domain (homologous to CheA), and response regulator domain but lack a homologous methyl-accepting domain (Fig. S1). Some HKRs (COP5-6/HKR1-2, and 8-10/HKR4-6) encode the response regulatory domain (homologous to CheY) and an adenylyl or guanylyl cyclase^24,26^. The light-switchable structural changes of COP5/HKR1 and COP6/HKR2 rhodopsin domains have been confirmed^27^. However, the control of guanylyl cyclase activity by light exposure has been confirmed in COP6/HKR2 but not any other HKRs^27^.

In *Chlamydomonas*, studies show that NH_4_^+^ chemotaxis depends on the presence of calcium ions and occurs optimally at an extracellular concentration of 10^_-3_^ M Ca^2+28,29^. However, how extracellular Ca^2+^ regulates the cilia-level response and what proteins are involved in this regulation is unknown. Calcium is commonly involved in chemotaxis signaling transduction, including in mammalian sperm^30^. For example, in mammalian sperm, a weakly voltage-dependent calcium channel CatSper, causes calcium influx into the cell when activated, switching the ciliary beat pattern to the symmetric waveform^3^. *Chlamydomonas* similarly has a voltage-gated calcium channel, CAV2, localized to the tip of the flagella responsible for changes in the ciliary beat pattern^31^.

*Chlamydomonas* has four types of ciliary beat patterns elicited by various environmental stimuli, which could be modulated during chemotaxis. Here, we categorize the ciliary beat patterns of *Chlamydomonas* as follows (Fig. 1):

**Figure 1.**
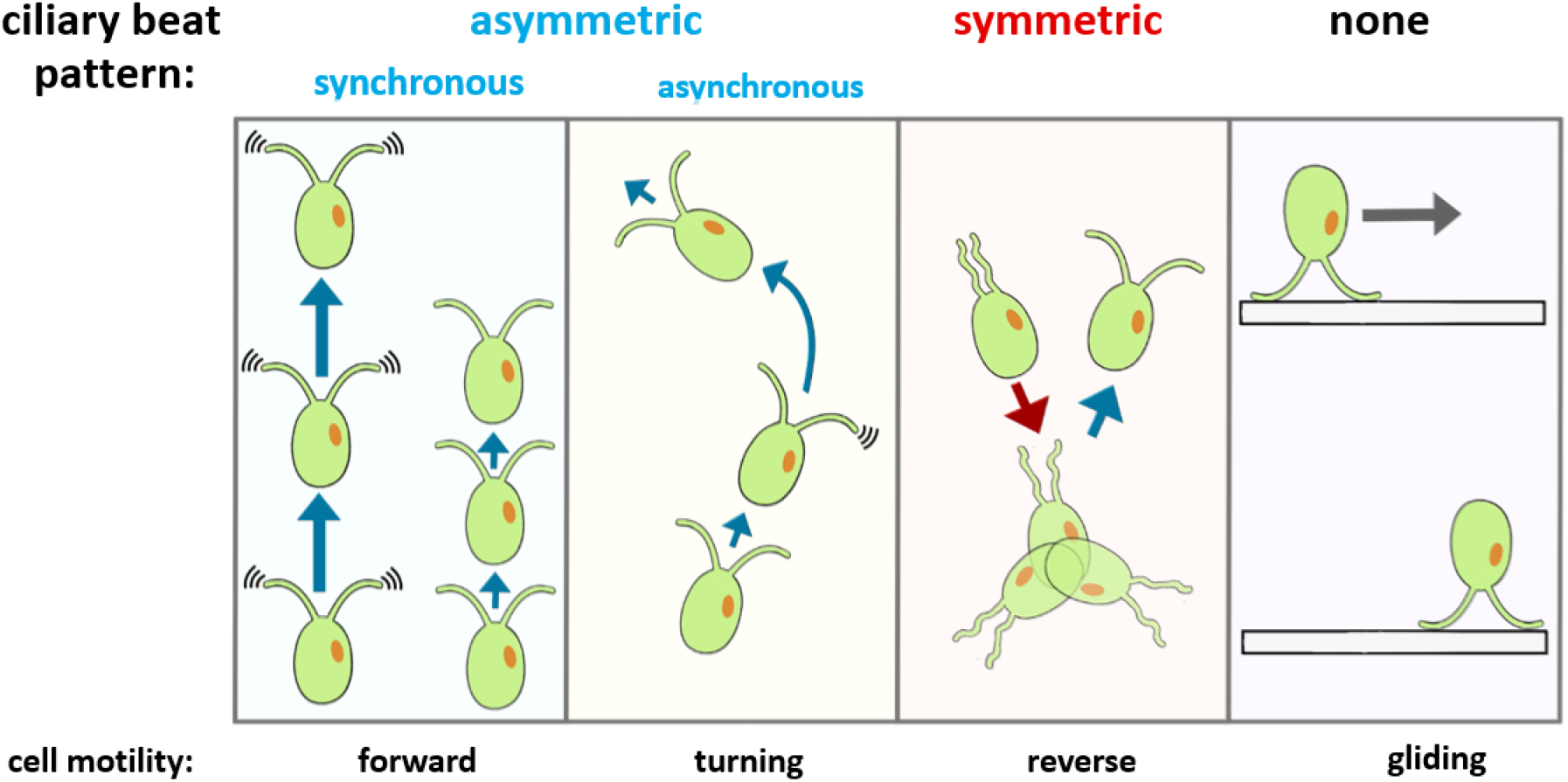
Four types of ciliary beat patterns: synchronous asymmetric waveform, asynchronous asymmetric waveform, symmetric waveform, and gliding in *Chlamydomonas*. *Chlamydomonas* has two general types of ciliary beats: the asymmetric waveform, pulling the cells forward (cilia move ahead of the cell), and the symmetric waveform, pushing the cells in reverse (cilia are moving behind the cell). The asymmetric waveform can be synchronous, making cells move straight at a speed based on beat frequency, or asynchronous, making cells turn due to unbalanced cilia propulsion. After the symmetric waveform, the cilia are uncoordinated for a short duration (less than a second) before switching back to the asymmetric waveform. Gliding occurs when cilia adhere to the surface of a material, and then the leading cilia pull the cell across the surface^*39*^.

1. Asymmetric and synchronous waveform, which is responsible for baseline movement, cell swimming speed, and photokinesis^32,33^.
2. Asymmetric and asynchronous waveform, which causes a turn in cell swimming angle, is responsible for phototactic steering^34,35^).
3. Symmetric waveform, a transient, spontaneously occurring waveform in wild-type strains, which causes a cell to swim in a reverse direction. This waveform is involved in the responses of photoshock^31^ mechanoshock^36^, and thermotaxis^37^.
4. Gliding, where the cilia adhere to and pull the cell along a surface. This movement can still occur in strains with disrupted ciliary beat, such as the paralyzed *pf18* strain^38^.

Based on previous chemotaxis studies in three domains of life and the homology of HKR proteins to sensor histidine kinases, we speculated that *Chlamydomonas* combines the signaling pathways of bacteria, archaea (histidine kinase sensor) and eukaryotes (guanylyl /adenylyl cyclase signaling, voltage-gated Ca^2+^ channel) to regulate cilia during ammonium chemotaxis (Table S1). Here, we report the results of our investigation, in which we identified the ciliary beat pattern used to steer up the gradient, signal transducer, and upstream signaling element responsible for ammonium chemotaxis in *Chlamydomonas*.

## Results

### Development of a medium-throughput chemotaxis assay for *Chlamydomonas*

We sought to identify *Chlamydomonas* strains deficient in ammonium (NH_4_^+^) chemotaxis. To this end, we designed and implemented a new Chemotaxis Lane Assay (CLA) (Fig. 2A). The CLA uses a 3D-printed reusable plate to determine the Chemotaxis Index based on changes in cell density. The CLA plate has two chemical reservoirs and 34 individual parallel lanes. The CLA plate is a flow-free system, relying on passive diffusion of the chosen chemical through permeable agarose hydrogel reservoirs to form a gradient in each lane. We first characterized the gradient formation in the CLA plate (Supplement Methods) using bromophenol blue (BPB), a dye molecule used to characterize diffusivity (Fig. 2B) ^40,41^. Assuming the chemical source acts as an infinite reservoir of constant concentration, the diffusion coefficient of BPB can be estimated by fitting the experimental data to the solution of Fick’s second law of diffusion (Eq. S1, Fig. S2C). The experimental diffusion coefficient was found to be 7.497 × 10^−5^ cm^2^ s^−1^, in concurrence with the reported diffusion coefficient of BPB (Fig. S2D).

**Figure 2.**
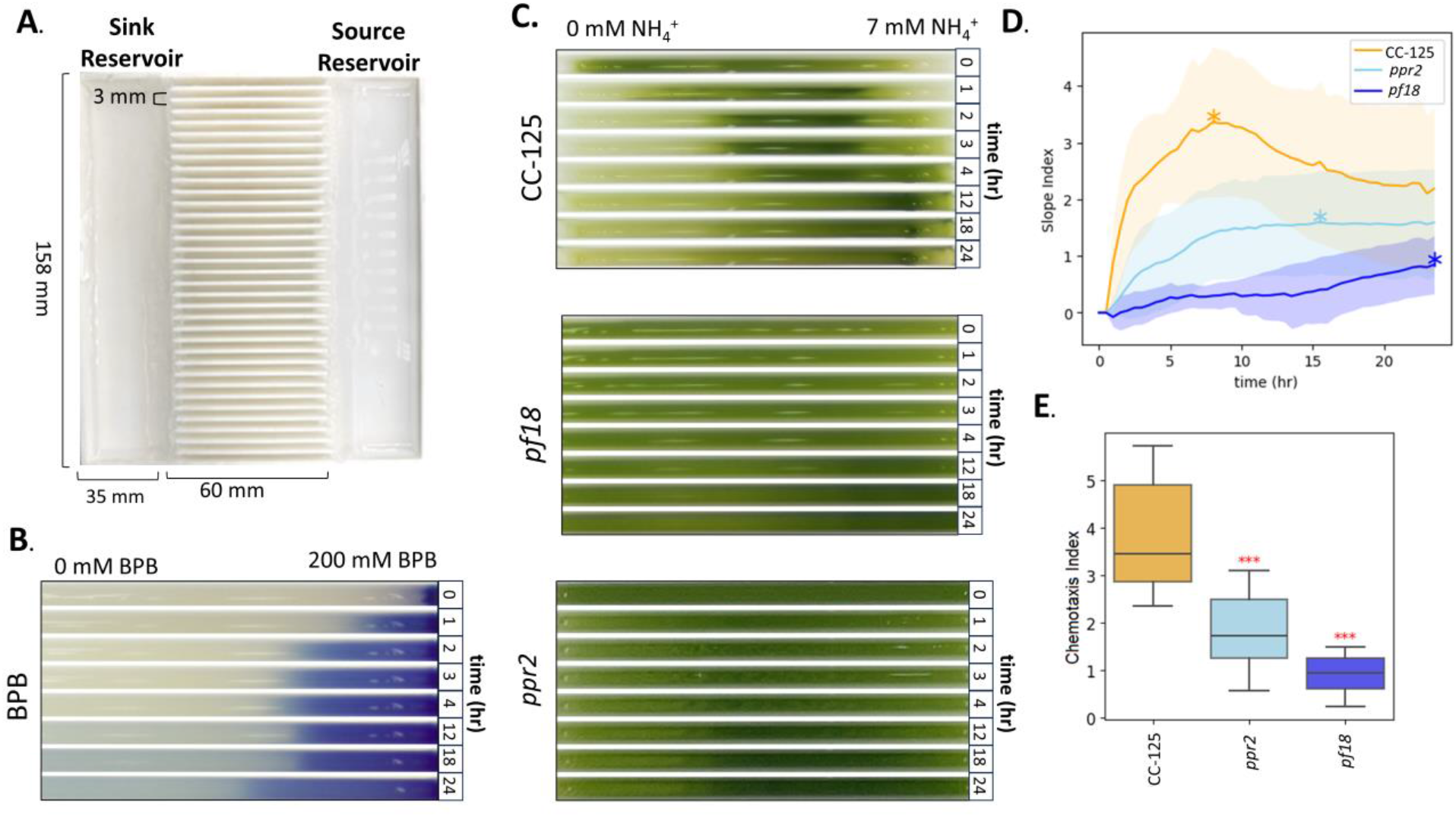
Identifying the *ppr2* strain (mutation in the *CAV2* gene encoding voltage-gated Ca_2_^+^ channel) as an ammonium chemotaxis-incompetent strain. **(A) Design and dimensions of the 3D-printed Chemotaxis Lane Assay (CLA) plate**. The plate has 34 lanes of 3 mm (width) x 60 mm (length) each. To use the plate, the source reservoir is set up with an agarose barrier containing 7 mM NH_4_^+^ (TAP+N) and loaded with TAP+N medium. The sink reservoir is set up with an agarose barrier containing 0 mM NH_4_^+^ (TAP-N) and loaded with TAP-N medium. **(B) Time-dependent chemical diffusion of BPB**. Bromophenol-blue (BPB) diffusion across a single lane at *t* = 0, 1, 2, 4, 12, 18, 24 hrs. The source reservoir set up with an agarose barrier containing 200 mM BPB and loaded with 200 mM BPB solution. The sink reservoir was set up and loaded with an agarose containing deionized water, and each lane was filled with deionized water. **(C) Chemotactic response of one lane of CC-125 (wild-type), *pf18* (paralyzed cilia mutant), and *ppr2* (*CAV2* gene mutant)**. The images show the chemotactic response of CC-125 (top), *pf18* (middle) and *ppr2* (bottom) at *t* = 0, 1, 2, 4, 12, 18, 24 hrs. **(D) Biomass density accumulation over time**. To characterize chemotaxis, the percent change from mean (PCM) of the pixel intensity after greyscale conversion (0-255) is calculated for each lane as a function of position and time (Fig. S3). The Slope Index is found by fitting the slope of the PCM across the 60 mm lane length, resulting in Slope Index for each time point. (*) represents the maximal Slope Index time of each strain. **(E) Comparison of Chemotaxis Indices**. For each replicate, the maximal Slope Index is then taken as the Chemotaxis Index and plotted as a boxplot. Significance was determined using a Student’s t-test where ***, p<0.001.

Next, we calculated the CLA’s Z-factor, a dimensionless statistical representation of the signal separation between the positive and negative controls, with values ranging from 1 (an ideal assay) to 0 or less (indicating no signal separation)^42^. We obtained Z’ = 0.69, categorizing the CLA as an “excellent” screening assay (Table S2, Fig. S2B). Additionally, the effect of lane position and day of the assay on the Chemotaxis Indices was examined to determine the replicability of the CLA (Fig. S2B). The observed positional variation is <8%, while the day-by-day effects are more considerable (<35%). This suggested that the Chemotaxis Indices may be higher or lower on some days, potentially due to initial cell density or metabolic state differences between trials. Considering this variation, all CLA experiments reported here were performed with replicates on at least three separate days and a positive control on each plate to increase statistical confidence in the resulting indices. Consequently, we classify the CLA as a medium throughput assay.

### The *Chlamydomonas* strain *ppr2* lacking the Ca^2+^-voltage gated channel subunit CAV2 is deficient in ammonium chemotaxis

When chemotaxis of the wild-type *Chlamydomonas* strain CC-125 was assayed with NH_4_^+^ in the CLA, alga accumulated at the NH_4_^+^ source and decreased at the sink, overlapping with the formation of the chemical gradient (Fig 2C). The Slope Index, a metric representing algal accumlation over time, obtained by taking the maximal slope of the percent change from the mean (PCM, Supplement Methods) of each lane, peaked around 6 hours for CC-125 (Fig. 2D). The paralyzed strain *pf18* showed a significant reduction in the chemotaxis response compared to the wild-type (Fig. 2D)^43,44^. Using the CLA, we found that *ppr2*, a strain that is phototaxis-competent but incapable of switching from the asymmetric to the symmetric waveform due to a mutation in the voltage-gated Ca^2+^ channel subunit CAV2^45^, exhibited reduced chemotaxis (Fig. 2E). The observed deficiency of *ppr2* suggested that mediation of waveform switching by the voltage-gated Ca^2+^ channel plays a crucial role in ammonium chemotaxis.

### Detection and characterization of cilia beat patterns in the *Chlamydomonas* cell population

To investigate the role of the symmetric waveform in ammonium chemotaxis, we characterized the relationship between switching ciliary beat patterns and cell motility (swimming speed and angle) during photoshock (Supplement Methods). Photoshock causes a population of cells to temporarily switch to the symmetric waveform, facilitating characterization of the symmetric waveform. By using single-cell tracking results from the open-source TrackMate plugin of ImageJ to identify changes in cell swimming speed and swimming angle, we confirmed that *randomized* swimming angle changes occur as a cell recovery from the symmetric to the asymmetric waveform, which occurs approximately 0.5 seconds after light flash illumination (Fig. S4, Fig. S5). This supports that switching to the symmetric waveform randomly changes a cell’s swimming trajectory, aligning with previous studies^46-48^.

Based on the observed changes in speed and angle, we developed a custom algorithm to automatically detect and quantify when individual cells switch from asymmetric to symmetric and back to asymmetric waveform (Fig. S6, Movie S1)^49,50^. *Chlamydomonas* cilia will spontaneously switch to the symmetric waveform when calcium is present in the assay media, even in the absence of an apparent external stimulus (i.e. photoshock, mechanoshock)^51^. To determine the incidences of this spontaneous waveform switching, our algorithm was applied to cells swimming in homogeneous (non-gradient) ammonium conditions, and the number of symmetric waveform incidence per track per second found. Subsequently, we found that the wild-type CC-125 strain spontaneously switches to the symmetric waveform 0.085 ± 0.032 times per second. In comparison to CC-125, the CAV2 deficient strain *ppr2* has a 1.6-fold decrease in incidences of switching to the symmetric waveform (Fig. S7).

### Calcium is required for ammonium chemotaxis signaling transduction

Due to our finding on CAV2, we wanted to examine the role of calcium in chemotaxis signaling transduction in the context of the ciliary beat pattern. To do so, we analyzed the relationship between cell motility and calcium availability in the media. Namely, the CLA (Fig. 2), photoshock assay (Fig. S5, Movie S1), phototaxis assay (Fig. S8), and homogenous (non-gradient) swimming assay (Supplement Methods) were conducted with and without the calcium chelator EGTA (ethylene glycol-bis(β-aminoethyl ether)-N,N,N′,N′-tetra acetic acid) in the assay media. Adding EGTA in the media decreased alga responsiveness across all motility assays (Fig. 4).

**Figure 4.**
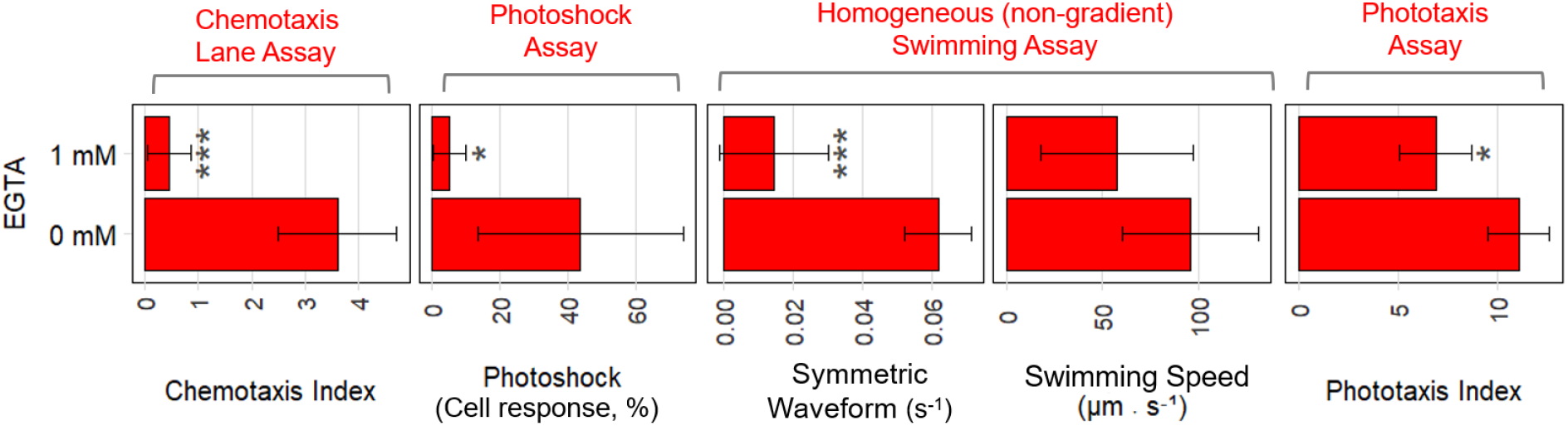
*Chlamydomonas* ammonium chemotaxis relies on the calcium-dependent symmetric waveform. The effect of EGTA, a chelator of Ca^2+^, on the motility of CC-125 (wild-type) was analyzed. Namely, the CLA, photoshock assay, homogenous (non-gradient) swimming assay, and phototaxis assay were conducted with and without EGTA. Two metrics were obtained from the homogenous (non-gradient) swimming assay: incidences of the symmetric waveform (Fig. S6), and swimming speed. Each bar represents the mean with the standard deviation of three independent experiments. Significance determined using a Student’s t-test, *** p<0.001, **p<0.01, *p<0.1, compared to 0 mM EGTA. EGTA was added to the assay medium ∼10 min before each experiment.

The decrease in motility in the presence of EGTA was most pronounced (***p<0.001) in the chemotactic index and incidences of the symmetric waveform. In *Chlamydomonas*, high intracellular calcium concentrations (10^−4^ M Ca^2+^) cause switching from asymmetric to symmetric waveform^52,53^. Therefore, the pronounced decrease in chemotaxis is likely due to the symmetric waveform being more sensitive to reduced calcium availability compared to the asynchronous and synchronous asymmetric waveforms, as demonstrated by the comparatively robust phototaxis and swimming speed. The results obtained in this analysis support that ammonium chemotaxis requires calcium for signaling transduction and uses different signaling transduction from phototaxis.

### *Chlamydomonas* strains lacking the *COP5* gene are incapable of sensing ammonium gradients

We revealed that *Chlamydomonas* requires the ability to perform waveform switching to bias its movement during ammonium chemotaxis and that calcium is involved in signaling transduction. A fundamental question remains: What molecule (protein) senses the ammonium gradient?

To answer this, we screened mutant strains deficient in one or more types of ciliary beat patterns as well as strains with knockouts of select members of the COP/HKR1 gene family (including two independent *COP5* gene-knockout mutants ΔCOP5-E6 and ΔCOP5.20-B2, generated from separate parental strains (CC-125 and CC-3043, respectively) by CRISPR technology^24^) with the CLA. We also analyzed the photoshock response, phototaxis, and swimming speed of these strains to determine the relationship between the four ciliary beat patterns. A hierarchical dendrogram is used to illustrate the relationship between cilia movement and the chemotactic response of the selected strains (Fig. 5A, Table S3).

**Figure 5.**
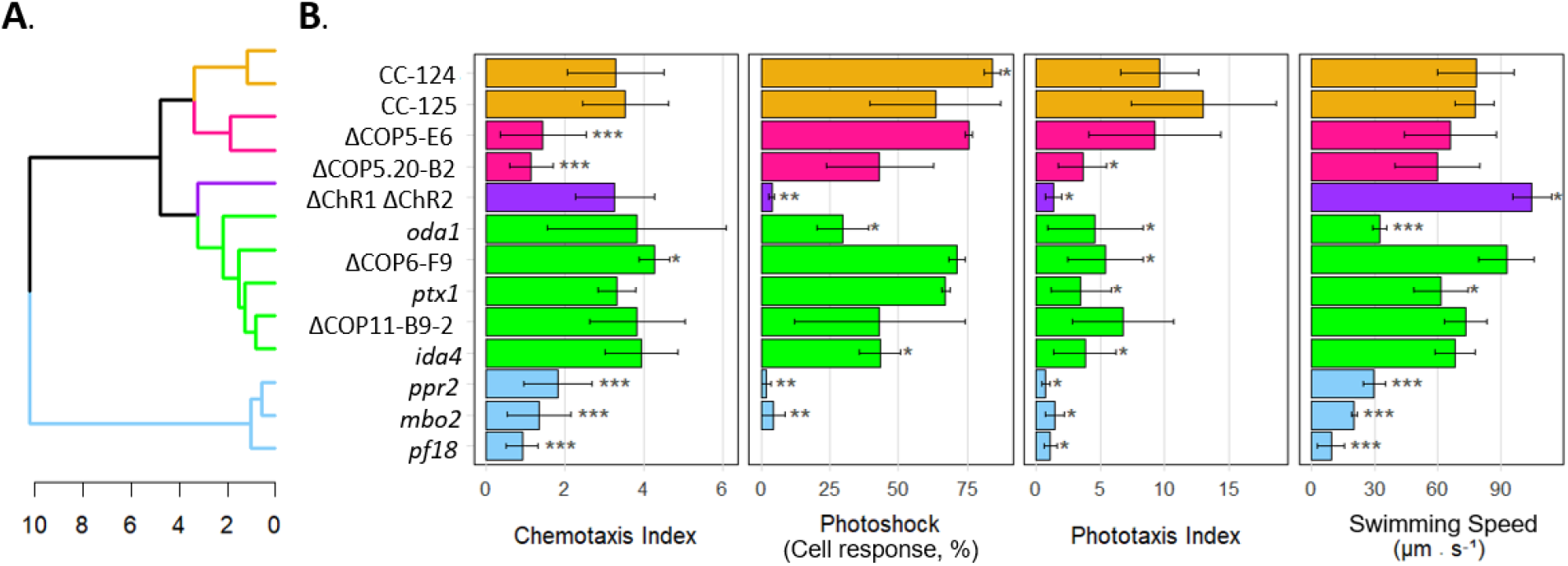
Identification of *COP5* knockout strains as an ammonium chemotaxis-incompetent strain. **(A) Clustering of *Chlamydomonas* strains based on the swimming phenotypes**. The dendrogram is colored by the k-mean cluster of the phenotypic averages and constructed using the hierarchal Ward linkage method^61^. **(B) Results of CLA, photoshock assay, phototaxis assay, and homogenous (non-gradient) swimming assay**. Each bar represents the mean with the standard deviation of a minimum of three independent experiments. P-values were calculated for each strain compared to CC-125 (wild-type), using a Student’s t-test, *** p<0.001, **p<0.01, *p<0.1. Notice that COP5 knockout strains show significantly lower Chemotaxis Index but normal values in other swimming phenotypes compared to CC-125.

The first group to examine is phototaxis mutants. *ptx1* has reduced phototaxis due to loss of control of cilia dominance required for control of the asynchronous asymmetric waveform due to a mutation in an unidentified gene^35^. This strain shows normal photoshock but reduced phototaxis (Fig. 5B). The mutant ΔChR1/ΔChR2 double knockout strain has a disruption in the two phototaxis receptors, leading to both reduced photoshock and phototaxis^54,55^. The CLA shows both strains maintains normal chemotaxis, demonstrating the separation of the known phototaxis signaling pathways and chemotaxis.

The strain *oda1* and *ida4* have defects in the outer and inner dynein arms, respectively, reducing ciliary beat frequency^5^. We found that *oda1* has a 58.3% reduction in swimming speed. Despite this reduction in baseline speed, *oda1* had normal ammonium chemotaxis, supporting that neither speed-based orthotaxis nor orthokinesis is involved in ammonium chemotaxis. *mbo2* is missing a coiled-coiled cilia protein and constantly expresses the symmetric waveform, making it deficient in photoshock and phototaxis^56,57^. Both *mbo2* (asymmetric waveform deficient) and *ppr2* (symmetric waveform deficient) have reductions in speed comparable to *oda1*, but unlike *oda1*, these strains consequently lose the chemotaxis response. Note that under our initial phototaxis assay conditions using white light, *ppr2* showed diminished phototaxis. When we performed the assay replicating prior studies using green light and incubation in a phototaxis assay solution^58,59,^ the phototaxis response was restored, matching previous reports (Fig. S9)^58^. The strain *pf18* which has lost all motility except for the gliding response does not exhibit ammonium chemotaxis, indicating gliding is not responsible for chemotactic steering^60^.

Among the strains assayed, ΔCOP5s were the unique exception. These strains exhibited a largely reduced chemotaxis response, yet the photoshock, phototaxis, and swimming speed are not significantly different from the wild-type strain. This suggests that deficient chemotaxis in ΔCOP5s is not due to an alteration of the ciliary beat pattern, as with *mbo2* and *ppr2*, but an upstream disruption of the ammonium gradient sensing mechanism. Strains with disruptions in the *COP5* gene family, *COP6* and C*OP11*^24^, exhibited wild-type like chemotaxis responses (Fig. 5). This indicated that the chemotaxis deficiency observed in ΔCOP5s is not a shared phenotype of the *COP* gene family.

### Wild-type *Chlamydomonas* strain, but not *COP5* knockout strain, biases waveform switching in an ammonium gradient

We found that ΔCOP5 strains have normal motility responses, except for chemotaxis, indicating COP5 is an upstream signaling element of ammonium chemotaxis (Fig. 5). Due to the identification of a mutant with a defect in upstream signaling, rather that cell steering, it can now be examined whether migration towards ammonium is a kinesis or taxis-like response. During a kinesis response cells change baseline motility in response to the *concentration* of a chemical, resulting in a passive accumulation. During a taxis response, cells bias motility in response to sensing the *gradient* steepness either temporally or spatially, resulting in an active and directed accumulation. Prior work indicates that *Chlamydomonas* is capable of performing both chemokinesis (accumulation due to biasing speed in response to sodium valerate) and chemotaxis (accumulation in the absence of a speed change in response to cobalt chloride and l-arginine) ^62^.

To determine if the ammonium response is kinesis-like, we tested the motility of CC-125 in varying homogenous ammonium concentrations. This experiment found that varying the ammonium concentration caused no statistically significant difference in any tested motility phenotype, including incidences of the symmetric waveform, indicating that the ammonium responses is not kinesis-like (Fig. S10).

After we found that the wild-type strain did not have a kinesis-like response, we tested if the ammonium response is taxis-like by examining ΔCOP5-E6 and CC-125 in an ammonium gradient. We performed a chemotaxis microscope assay with a chambered cover glass to form an ammonium gradient (Fig. 6, Movie S3). Cells in the ammonium gradient show visible migration towards the ammonium source during the assay in CC-125 but not ΔCOP5-E6 (Fig. 6B, Movie S2). To determine if the symmetric waveform is correlated to cell swimming orientation in a gradient, we analyzed the relative incidences of the symmetric waveform for cells swimming toward the source (7 mM NH_4_^+^) or toward the sink (0 mM NH_4_^+^), in the chambered cover glass (Fig. 6). In this analysis, if *Chlamydomonas* biases the incidences of the symmetric waveform, the ratio of the relative symmetric and asymmetric waveform incidences should be different between the cells moving toward the source and those toward the sink.

**Figure 6.**
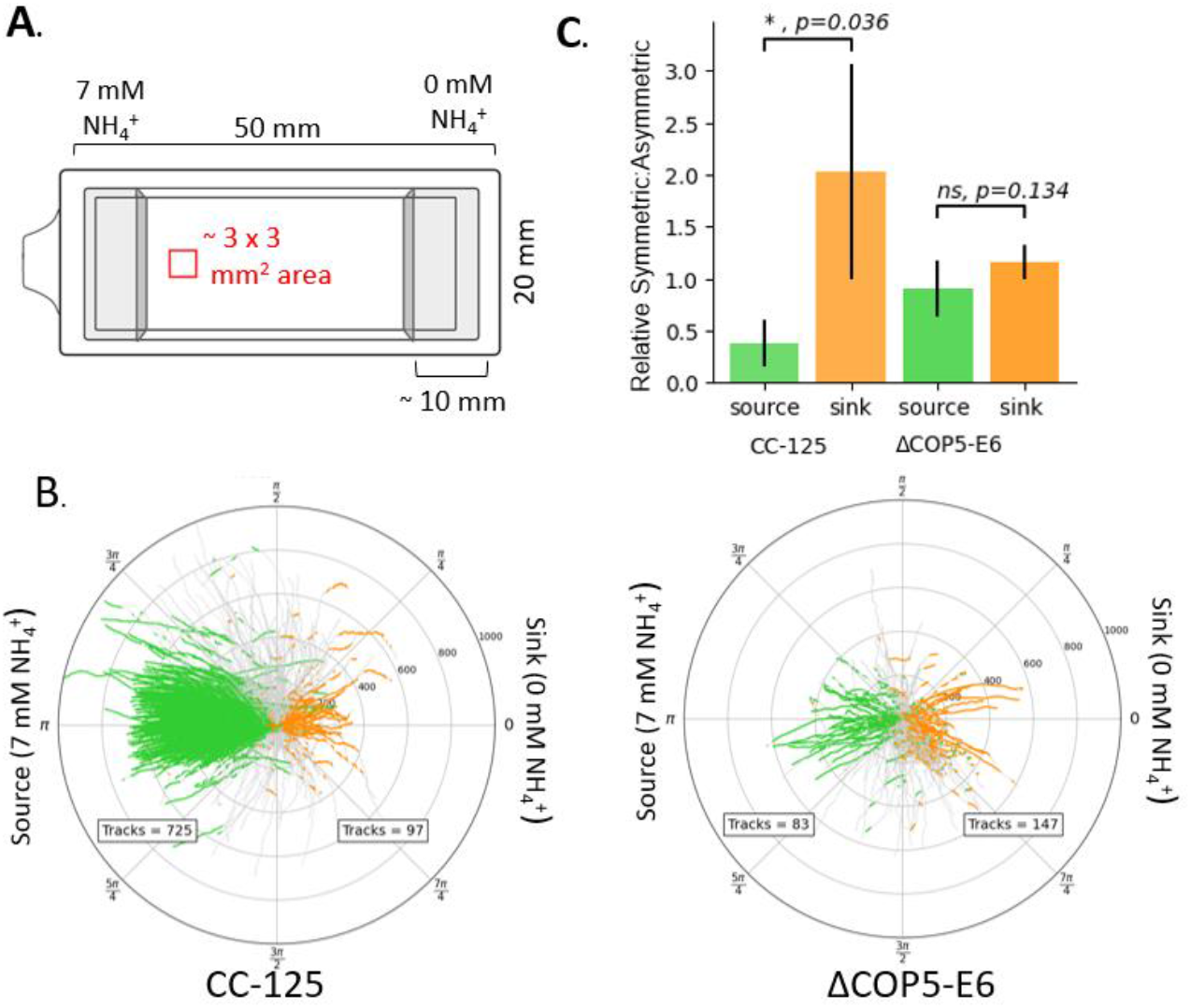
Wild-type *Chlamydomonas* strain but not *COP5* knockout strain bias switching to the symmetric waveform in an ammonium gradient. Timelapse videos of wild-type (CC-125) and *COP5* knockout (ΔCOP5-E6) strains were tracked at 30-fps (frame per second) in a flow-free ammonium gradient (see Movie S2) taken at *t* = 60secs after initial exposure to the ammonium gradient, for 10 secs. Tracks with durations shorter than 4 secs are discarded. **(A) Schematic drawing of a top view of a chambered coverglass used in a chemotaxis microscope assay**. A 7 mM NH_4_^+^ (TAP) containing agarose block was placed on the left side of the chamber, and a 0 mM NH_4_^+^ (TAP-N) containing agarose block was placed on the right side. The red square indicates the approximate location where cells were tracked. **(B) Orientation angle analysis of single-cell trajectories**. Single-cell tracking of each strain was performed and the orientation angle relative to the source was determined. A green track segment indicates cell movement toward the source (7 mM NH_4_^+^). An orange segment indicates cell movement toward the sink (0 mM NH_4_^+^).**(C) Statistical analysis on relative incidences of the symmetric waveform**. The ratio of the relative symmetric waveform to asymmetric waveform in the tracks was calculated in the population of the cells that moved toward the source (green bars) and sink (orange bars), respectively (Supplement Methods). Each bar represents the mean of the “Relative Symmetric:Asymmetric” waveform ratio, and error bars represent the standard deviation (n=4). P values comparing the source and the sink for each strain are found using the Student’s t-test.

The analysis revealed that switching to the symmetric waveform is significantly lower when cells swam toward the source compared to when cells swam toward the sink in the wild-type strain CC-125 (Fig. 6). These result support that ammonium chemotaxis is taxis-like, as a biasing cell locomotion occurs in an ammonium gradient but not homogenous ammonium conditions. In comparison, ΔCOP5-E6 did not show a significant bias in ratio of the relative symmetric and asymmetric waveform incidences supporting that COP5 is an essential component to control the chemotaxis response of *Chlamydomonas* in an ammonium gradient.

## Discussion

In this study we reveal that the symmetric waveform is required to steer in an ammonium gradient in *Chlamydomonas*. Similarly to the “run and tumble” mechanism of bacterial chemotaxis, when *Chlamydomonas* switches from the symmetric to the asymmetric waveform, random direction changes occur (Fig. S4), biasing cell position based on the principle of a biased random walk^63,64^. In *Chlamydomonas*, switching from the asymmetric to symmetric waveform and back (causing random directional changes) occurs at a lower rate when swimming up the gradient, leading to the directed accumulation of cells at the chemical source (Fig. 7A). This is the first time the “run and tumble” steering strategy has been directly observed in response to a chemical gradient for ciliate green microalgae.

**Figure 7.**
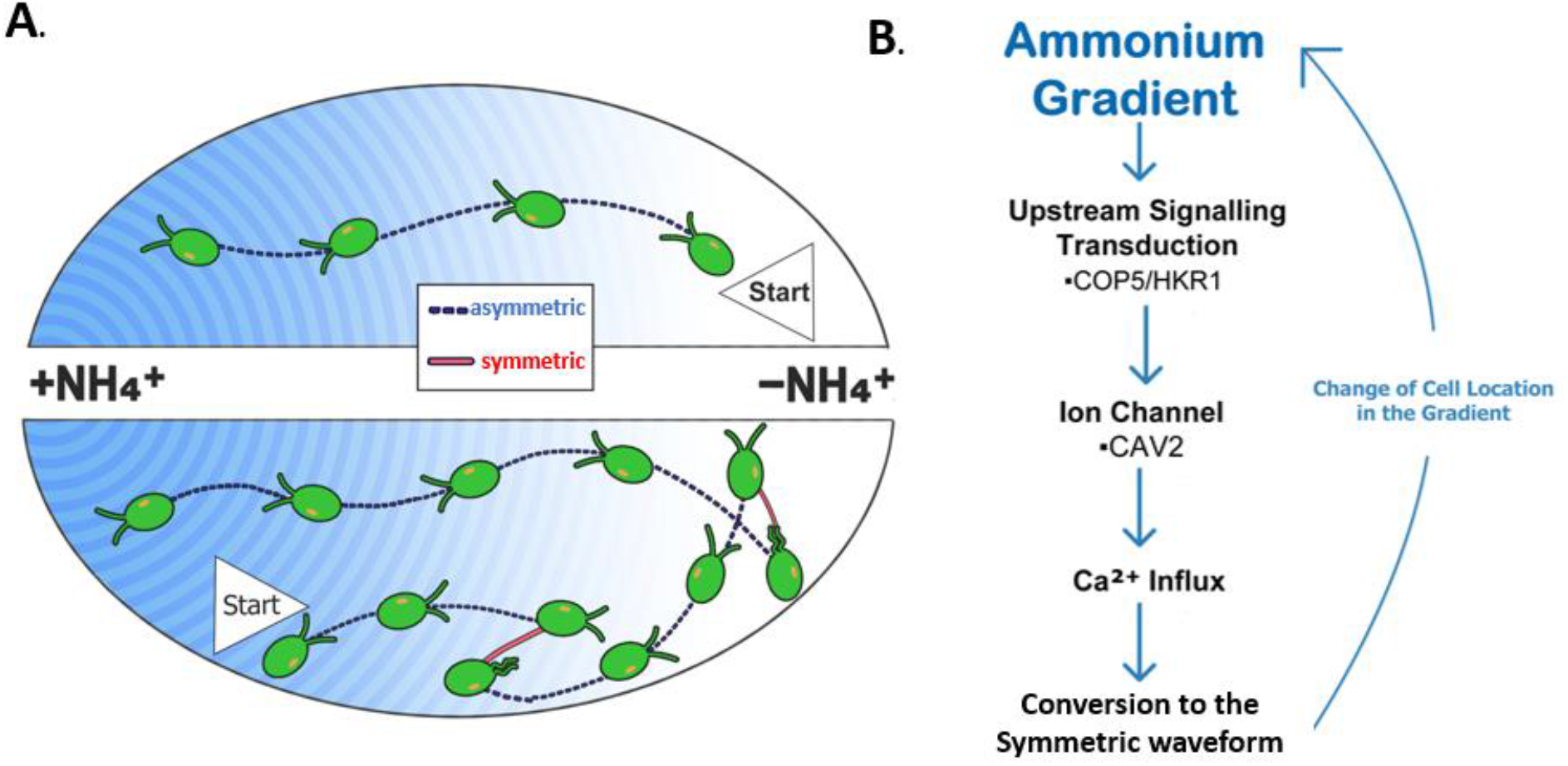
Ammonium chemotaxis steering mechanism and proposed signal transduction pathway. **(A) The steering mechanism for NH_4_^+^ chemotaxis**. During swimming, a cell switches ciliary beat patterns. When the ciliary beat pattern switches from the symmetric to asymmetric waveform, the cilia are briefly uncoordinated, leading to a random directional change. Waveform switching is lower when cells swim up an ammonium gradient, causing a biased random walk and net movement of the population toward the chemical source. **(B) The proposed signaling transduction pathway of ammonium chemotaxis identifies COP5/HKR1 as an upstream signaling element**. COP5/HKR1 senses the gradient and mediates the CAV2 (voltage-gated Ca^2+^ channel) activity downstream. Ca^2+^ influx triggers the switch of ciliary beat patterns.

Two proteins out of the 12 COP family members in *Chlamydomonas*, COP3, and COP4, more commonly known as Channelrhodopsin 1 and 2 (ChR1 and ChR2), have been identified as phototaxis light receptors (Fig. S1)^25^. However, the biological function of the remaining COP proteins, including ones that have a histidine kinase encoded in the C-terminal of the rhodopsin domain (the alternative name of these COP members is histidine kinase rhodopsin, HKR), has not been identified. We discovered that *Chlamydomonas* mutant strains lacking COP5/HKR1 are deficient in ammonium chemotaxis. In contrast, the strains lacking COP6/HKR2 and COP11/HKR7 maintain ammonium chemotaxis. This suggests that COP5/HKR1 may play a uniquely role in ammonium response among the COP/HKR protein family. While mRNA expression of all HKRs has been confirmed^26^, COP5/HKR1 is recognized as the most abundantly expressed HKR gene in the *Chlamydomonas* gene expression database^65^.

Here, we postulate the signaling transduction pathway of ammonium chemotaxis in *Chlamydomonas*, where signaling is regulated by COP5/HKR1 upstream and transduced with the voltage-gated Ca^2+^ channel, causing a calcium influx and leading to switching the ciliary beat pattern (Fig. 7B). There are likely several unidentified intermediate components in the chemotaxis signaling pathway. Specifically, how COP5/HKR1 interacts with ammonium at a biochemical level and transduces the chemotactic signal to the voltage-gated Ca^2+^ channel.

Several possibilities to consider include that while COP5 may directly bind to ammonium, it is also possible that COP5/HKR1 interacts with a separate receptor protein responsible for the initiation of chemoreception, such as one of the eight identified ammonium transport channels^66^. In bacterial and archaeal chemotaxis, the ligand binding domain of MCP proteins is responsible for chemoattractant binding. However, all *Chlamydomonas* HKRs lack the MCP domain, and no homologous proteins to the *E. Coli* MCP proteins are found in the *Chlamydomonas* genome^26,67^. Interestingly, COP5/HKR1 encodes an extra transmembrane domain (known as domain zero) in the rhodopsin domain, normally composed of the seven transmembrane domains^68^. The biochemical function of the domain zero of the HKRs is unknown but this domain may play a role in ammonium reception. At this point, further biochemical and genetic studies are needed to characterize the precise role that COP5/HKR1 plays in chemoreception.

How COP5/HKR1 transduces the signal to the voltage-gated calcium channel, CAV2, also invites further investigation. COP5/HKR1 may directly bind to and interact with another downstream signaling protein, as is predicted to be the case with mammalian sperm calcium channel CatSper^3^. To this point, COP5/HKR1 encodes a sterile alpha motif domain at the C-terminal end that is known to be involved in protein-protein interactions^69^. Due to the presence of the guanylyl cyclase domain, it is more likely that cGMP functions as a secondary messenger to transduce the chemotactic signal. However, the cyclase activity of COP5/HKR1 has not yet been proven, and its cyclase domain lacks homology to the functional COP6 guanylyl cyclase in key residues^27^. Another component to consider is that COP5/HKR1 may act similarly to the phototaxis receptors ChR1 and ChR2 by directly or indirectly initiating ion influx into the cell, causing membrane depolarization, and activating downstream calcium influx through the voltage-gated CAV2^55^. The calcium sensor protein that would sense this influx and transmit the signal to the cilia and induce symmetric waveform conversion is currently unidentified.

Lastly, *Chlamydomonas* exhibits chemotaxis towards six other attractants and repellents: bicarbonate, sugars (maltose, sucrose, xylose, and mannitol), cobalt (II) chloride, manganese (II) sulfate, l-arginine and tryptone^11,62,70,71^. It is possible that the other COP proteins, COP6/HKR2-COP12/HKR8, could play a role in the chemoreception of these species, similarly to COP5/HKR1.

## Materials and methods

### Algal strains and cultivation

*Chlamydomonas* strains (Table S3) were grown in 100 ml Erlenmeyer flasks or 15 ml polypropylene round bottomed tubes in Tris-acetate phosphate (TAP) to a density of 10^6^ cells/ml^72^. Cultures grown on G10 Gyrotory Shaker at 110 rpm, under 30 photons·m^−2^·s^−1^ µmol lights, on a 12:12-hour (dark: light) cycle. All experiments started during the first 3 to 6 hours of the dark cycle and performed at room temperature. Cells were collected by centrifugation at 2100 rpm for 5 minutes, washed with Tris-acetate phosphate without the nitrogen (TAP-N) once, then re-suspended to a final concentration of 10^6^ cells · ml^−1^.

### Chemotaxis Lane Assay (CLA)

The CLA-Plate was fabricated with 34 independent parallel lanes, the outer two of which were not utilized for experimental purposes. 3D STL models were created using three-dimensional (3D) solid modeling software (SOLIDWORKS). The .stl files from SOLIDWORKS were converted to .gcodes formed for printing in the slicing software Simplify3D. CLA-Plates were printed using ABS (acrylonitrile butadiene styrene) filament on an Ender 3 Series 3D fused deposition modeling (FDM) printer. The CLA plate was disinfected with a 7.5% sodium hypochlorite solution between experiments.

To set up the CLA-plate to test for NH_4_^+^ chemotaxis, the sink and source reservoirs are filled with TAP-N (0 mM NH_4_^+^) or TAP (7 mM NH_4_^+^) respectively and separated from the lanes by two permeable agarose barriers each matching their respective reservoir solutions (1.5% agarose, ∼3-4 mm wide). Each lane is filled with 1 ml of algae suspended in TAP-N. The assay was performed in a Photobox and illuminated from the top of the box with homogenous white light (24 µmol m^−2^s^−1^). All sides of PhotoBox are covered with a reflective Mylar film (Mylar Diamond Film, TEXALAN). Experimental timelapse was taken at 30-minute intervals for 24 hours.

Images of the CLA-Plate taken every 30 minutes for 24 hours on a Raspberry Pi powered camera (12.3-megapixel Sony IMX477 sensor, 7.9mm diagonal image size). Images were imported into ImageJ imaging software^73^, aligned using the ImageJ plugin Linear Stack alignment with SIFT, converted to 8-bit, and inverted. Pixel intensity is recorded vertically across the 64 mm x 1.5 mm lane, at every time point, resulting in a dataset of pixel intensity as a function of time and position.

### Chemotaxis Microscope Assay

A chambered coverglass (Lab-Tek II Chambered #1.5 German Coverglass System) was used to observe chemotaxis under a microscope. The interior dimensions of the chamber are 10 mm (height) x 20 mm (width) x 50 mm (length). The right side of the chamber is filled with a 1.5% agarose hydrogel block loaded with 7 mM NH_4_^+^ TAP, and the left side is filled with a 1.5% agarose hydrogel block loaded with 0 mM NH_4_^+^ TAP, resulting in two agarose blocks of 10 mm (height) x 20 mm (width) x ∼ 10 mm (length). Sixty seconds after the chamber was loaded with algae, video timelapses were captured for ten seconds. Microscopy was performed with an Olympus IX83 Inverted Microscope equipped with ORCA-Fusion BT CMOS camera and a 4x/N.A.0.16 objective lens. Timelapse images were taken at 30 fps (frame per second) with 2×2 binning. A LED ring light (AmScope 144 LED Intensity-Adjustable Ring Light) was used to provide homogenous white light exposure to from the top of the chamber.

## Acknowledgement

We thank Dr. Pinfen Yang for providing *pf18*.

## Code availability

Code required to generate figures is available upon request

