## Supporting Information for "Membrane-bound Guanylyl Cyclase COP5/HKR1 changes ciliary beat pattern and biases cell steering during chemotaxis in *Chlamydomonas reinhardtii*"

### Supplemental Tables

**Table S1.** Mechanism of chemotaxis in model cells that utilize a motile organelle protruding from the cell surface

| Species | Kingdom | Domain | Motile organelle | Steering mechanism | Ligand receptor | Signaling pathway |
|---|---|---|---|---|---|---|
| *E. coli(1)* | Bacteria | Bacteria | Flagella | Run and tumble | MCP* | Histidine Kinase |
| *H. salinarum(2)* | Archaea | Archaea | Archaea | Forward and reverse | MCP * | Histidine Kinase |
| *P. lividus* (sperm)(3, 4) | Animalia | Eukaryota | Cilia | Relaxing helical path | Speract receptor | cGMP**, $vgCa^{2+***}$ |
| ***C. reinhardtii*** | Plantae | Eukaryota | Cilia | ***Unknown*** | ***Unknown*** | ***Unknown*** |

*MCP: methyl-accepting protein, **cGMP: Guanosine 3',5'-cyclic monophosphate, $vgCa^{2+}$: voltage gated $Ca^{2+}$ channel

**Table S2.** Z-Factors and analysis of positional and day-by-day effects of Chemotaxis Lane Assay

| **conditio*n*** | ***group*** | ***z'*** | ***Percent difference from mean (%)[a]*** | |
|---|---|---|---|---|
| | | | Max signal | Min signal |
| day | 1 | 0.985056 | 15.700 | -19.142 |
| | 2 | 0.751416 | 20.701 | 62.528 |
| | 3 | 0.911677 | -36.400 | -43.386 |
| position | Left (L) | 0.472473 | 6.351 | 1.417 |
| | Middle (M) | 0.750746 | 1.014 | 21.160 |
| | Right (R) | 0.673839 | -7.364 | -22.577 |
| pooled data | -- | 0.698692 | -- | -- |

a. Percent difference is defined as the difference in the mean of each individual group, to the pooled mean across all groups.

**Table S3.** Strains used in study

| Strain number | Strain name [gene, if known] | Parental strain | Phenotype | Mutant method | Source[a-c] | Reference |
|---|---|---|---|---|---|---|
| -- | *ppr2* [CAV2] | CC-125 | Loss of symmetric waveforms due mutation of α-subunit of voltage-gated $Ca_2^+$ channel | nit1 gene insertion | WAKA | (5) |
| -- | *pf18* [PF18] | --- | Paralyzed cilia. Almost no swimming motility. | -- | YANG | (6) |
| CC-124 | [NIT1 NIT2 AGG1] | ---- | Wild-type | ---- | CRC | (7) |
| CC-125 | [NIT1 NIT2 AGG1] | ---- | Wild-type | ----- | CRC | (7) |
| CC-2228 | *oda1* [DC2] | 137c+ | Missing outer dynein arm and the outer dynein arm docking complex, slow and jerky swimming | UV light mutagenesis | CRC | (8) |
| CC-2377 | *mbo2* | 137c | Loss of asymmetric waveforms | Chemical mutagenesis | CRC | (9) |
| CC-2670 | *ida4* [Dll1] | *oda1* | Missing inner dynein arm (a, c and d), slow swimming | Chemical mutagenesis | CRC | (8, 10, 11) |
| CC-2894 | *ptx1* | 137+ | Deficient in $Ca^{2+}$ dependent control of cilia dominance | UV light mutagenesis | CRC | (12) |
| CC-5428 | ΔCOP11-B9-2 [COP11] | CC-125 | Missing chlamyopsin, function unknown | CRISPR/Cas 9 | CRC | -- |
| CC-5431 | ΔCOP6-F9 [COP6] | CC-125 | Missing chlamyopsin, function unknown | CRISPR/Cas 9 | CRC | -- |
| CC-5437 | ΔCOP5.20-B2 [COP5] | CC-3403 | Missing chlamyopsin, | CRISPR/Cas 9 | CRC | (13) |

| | | | | | | |
|---|---|---|---|---|---|---|
| | | | function unknown | | | |
| CC-5448 | ΔCOP5-E6 [COP5] | CC-125 | Missing chlamyopsin, function unknown | CRISPR/Cas 9 | CRC | (13) |
| CC-5499 | ΔChR1 ΔChR2 [COP3, COP4] | CC-125 | Missing phototaxis light receptors | CRISPR/Cas 9 | CRC | (14) |

A. WAKA: Strain provided by Dr. Ken-ichi Wakabayashi
B. YANG: Strain provided by Dr. Pifen Yang
C. CRC: Strain obtained from the Chlamydomonas Research Center

### Supplement Figures

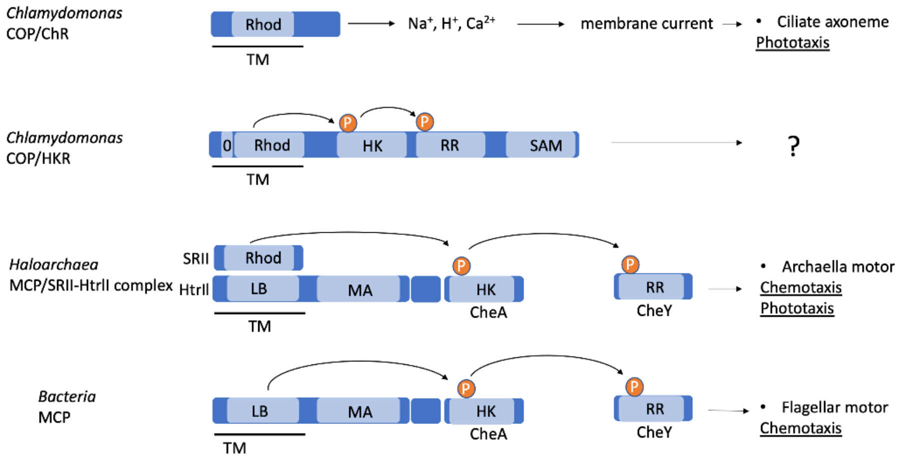


**Fig. S1. Sensor histidine kinase in bacteria and archaea, and *C. reinhardtii* chlamyopsin.** Sensor histidine kinase is the major chemoreceptor found in bacteria and archaea. Homologous proteins are found in fungi and plants as well. In bacteria and archaea, the "two-component system" composed of multiple proteins transduces the chemotactic signal (15). The signal initiated by the ligand binding domain (LB) is transferred through phosphorylation of the histidine kinase domain (HK), also known as CheA, and a response regular protein (RR), also known as CheY(16). The methyl-accepting domain (MA), where methylation/demethylation occurs, functions as a feedback regulator for the HK activity(17). In archaea, the chemoreceptor dual functions as a signal transducer of phototaxis by forming a complex with the photoreceptor rhodopsin (Rhod)(18). In *C. reinhardtii*, chlamyopsin 3 and 4 (COP3 and 4) encoding Rhod functions to transduce the phototaxis signal by importing ions into the cells. *C. reinhardtii* possesses homologous proteins of the histidine kinase receptor HKRs (histidine kinase rhodopsin). HKRs are the members of the COP family (19). Receptor histidine kinase in *C. reinhardtii* encodes Rhod followed by HK and RR (guanylyl cyclase) but not MA. They also encode an extra transmembrane domain (0) in the Rhod domain. COP5, 6, and 8 also encode the sterile alpha motif domain (SAM) that is known to be involved in protein-protein interactions (20). The biological function of the histidine kinase receptor in *C. reinhardtii* is unknown. TM: transmembrane region. The terminal recipient of the signal transduction of each protein (complex) is described with a bullet and the biological function is underlined.

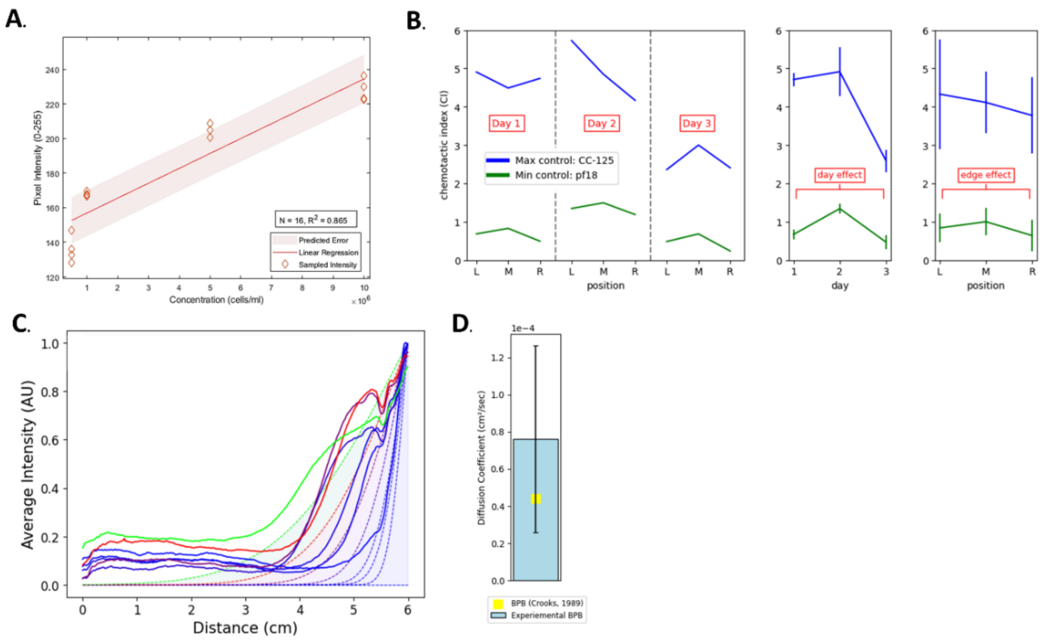


**Fig. S2. Determination of Chemotaxis Lane Assay (CLA) quality (A) Pixel intensity of a CLA-plate with *Chlamydomonas* suspensions of differing cell densities**. Individual lanes filled with 1 ml of algae at varying concentrations and imaged in a photobox using the same setup described previously(21). A linear relationship between cell density and pixel intensity was found. **(B) Variation in positional and day-by-day responses of CLA**. CLA was performed across three days using CC-125 as a positive control representing the maximal chemotactic response and *pf18* as a negative control representing the minimal chemotactic response, at three positions on the CLA-plate, L=Left, M=Middle, R=Right. Graphs to the left show the unaveraged results. Graphs to the right show the responses for CC-125 and *pf18* averaged by either day or position, corresponding to Table 2. **(C) Bromophenol blue (BPB) gradient formation.** The relative BPB concentration is represented as normalized intensity (AU), normalized after taking the average pixel intensity of the experimental BPB diffusion (N=28 lanes, solid line). Experimental intensity data is plotted against the curve-fitted solution of Fick's second law (dotted line**). (D) Experimental and reported diffusion coefficients.** The experimental diffusion coefficient, D, is found by curve fitting to Equation S1 and plotted against the previously reported BPB diffusion coefficient of 4.4 x $10^{-6}$ $cm^2$ $s^{-1}$ (22).

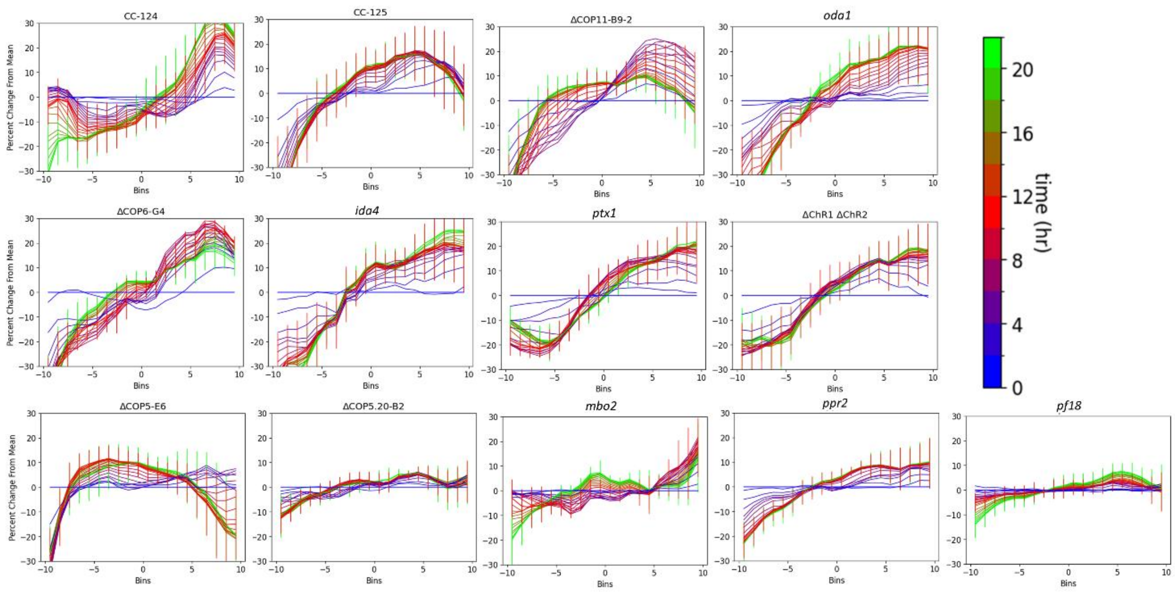


**Fig. S3. Average percent change from mean of Chemotaxis Lane Assay for each strain.** Percent Change from Mean (PCM) of all strains, shown as a function of time (indicated by the color of each line corresponding to a color-lookup table from 0 (blue) to 24 hours (green), and bin (corresponding to the lane length). The Y-axis of each graph shows the 60-millimeter lane is divided into 20 bins where bin "-10" is closest to the chemical sink, bin "+10" is closest to the chemical source, and "0" is the middle of the lane. An increase in the PCM on the right-hand side of each graph indicates accumulation at the chemical source, and on the left, a decrease indicates migration away from the chemical sink. Error bars shown in $\mathbf{t} = 0$, 12, 24 hrs. PCM shown for at least 3 replicates.

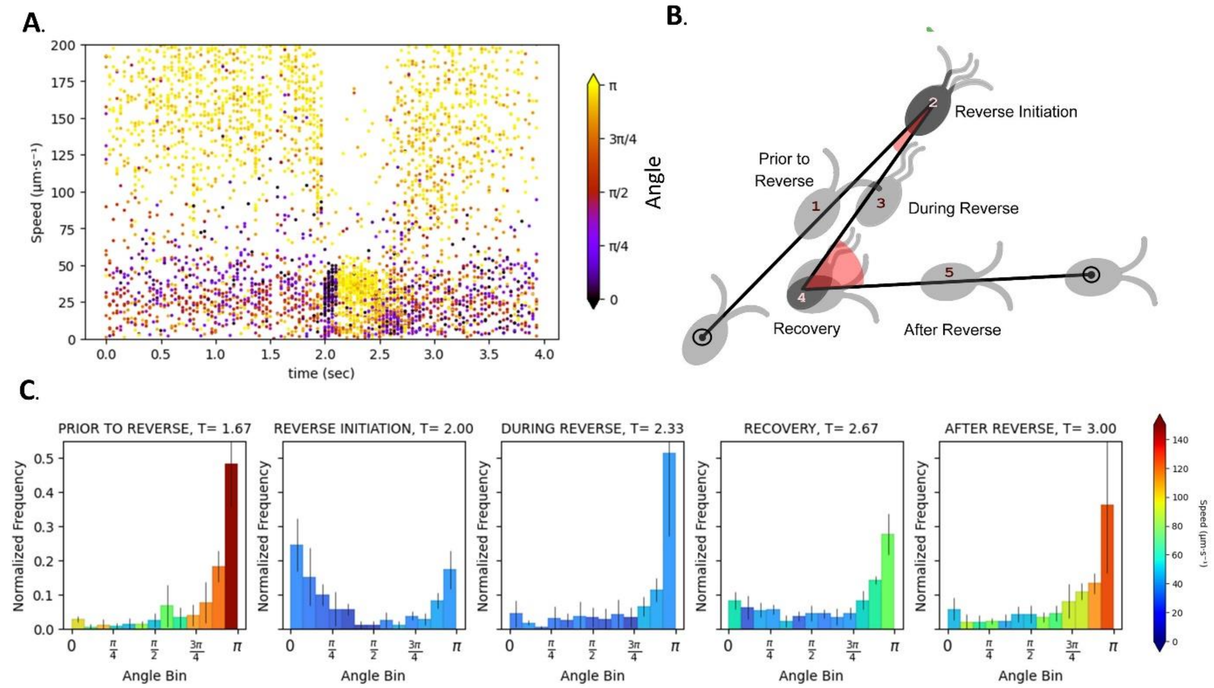


**Fig. S4. Characterization of symmetric waveform and recovery during photoshock. (A) One CC-125 trial shows swimming angle and speed changes during the photoshock response**. Each plotted point represents the speed of one cell at one time-point, colored by cell swimming angle from 0 (backward, indigo) to π (straight, yellow). Light flash illumination occurs at t = 2.0 sec. (**B) Schematic description of each stage of symmetric waveform**. Five stages, classified as "prior to reverse", "reverse initiation", "during reverse", "recovery" and "after reverse" with distinct swimming angle and speed distributions occurs as shown in part C. **(C) Histograms of average speed and swimming angle distribution of cell populations during photoshock.** For three independent photoshock experiments of CC-125, swimming angle distribution was analyzed at 5-time points, t=1.67, 2.00, 2.33, 2.67, and 3.00 sec. Each histogram shows the normalized swimming angle distribution of the population within a 0.166 sec interval following each respective time point. For each time interval, the swimming angles of all trajectories were divided into 13 bins ranging from 0 to π, resulting in a histogram. Before averaging, each histogram bin was normalized by dividing by the total swimming angle count. Each histogram bar represents the average normalized swimming angle frequency of three photoshock experiments and error bars show standard deviation. The color of each bar corresponds to the average trajectory speed in each swimming angle bin. Each histogram timepoint corresponds to the Symmetric waveform stages, as shown in the panel B.

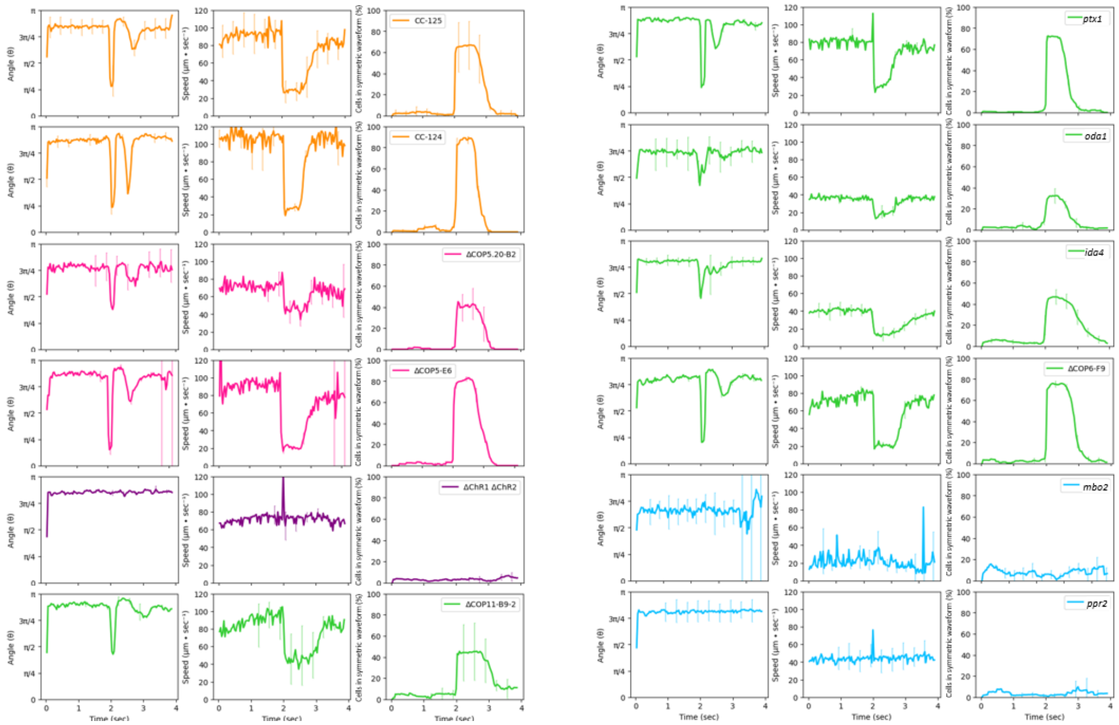


**Fig. S5. Photoshock response curves and changes in speed and angle.** Photoshock assay was performed with a light shock stimulus applied at t = 2 sec. The average speed, angle, and percentage of cells with the symmetric waveform across three independent photoshock assays is calculated for a 4-second time interval. The swimming angle at each point of the cell trajectory is found by taking the angle at each time (t), as t – 0.133 sec, t, and t + 0.133 sec, where 0 is a swimming angle with a complete reverse and π is a straight swimming angle. The number of trajectories in symmetric waveform at each time point (identified with custom Python algorithm, Fig. S6) was divided by the total number of tracked cells at each time point to find the percentage of cells in symmetric waveform. The plotted line represents the average of each photoshock metric, and the error bars represent the standard deviation. Each strain is colored by phenotypic cluster as identified in Fig 5.

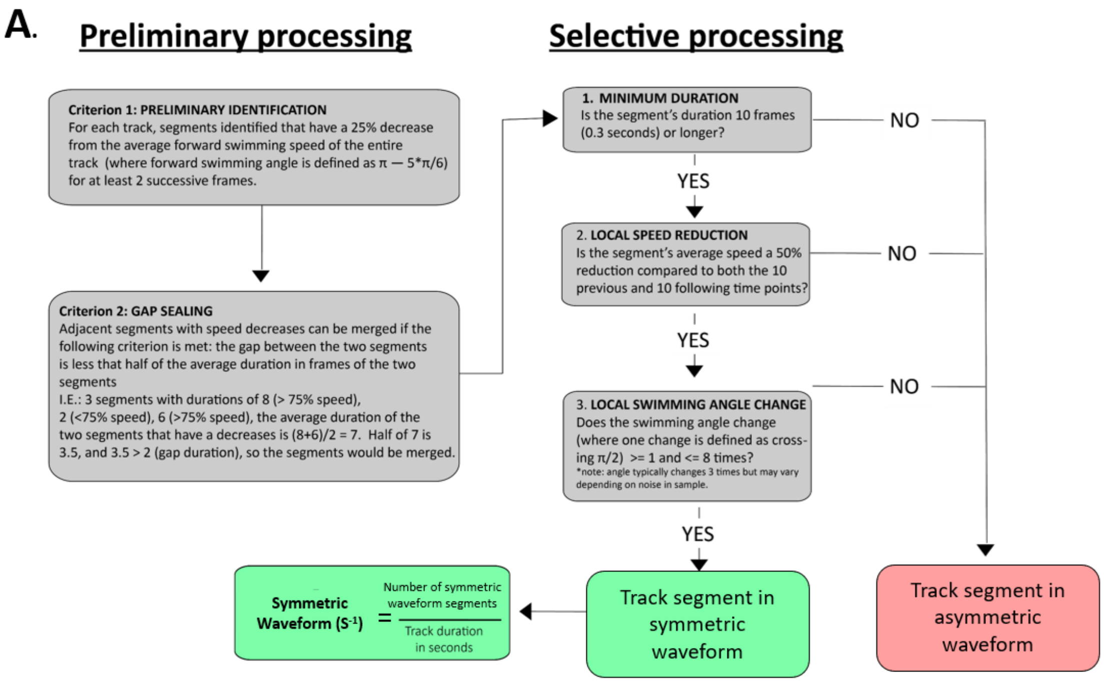


**Fig. S6. Algorithm to identify and quantify symmetric waveform in single-cell trajectories.** The flow chart details logic of a custom Python code used to identify reverse-recovery swimming. Code applied to analyze three separate assays: the photoshock assay (to calculate the percentage of cells that respond to a photoshock stimuli; Fig. 4, Fig. 5, Fig. S5, Fig. S10), homogeneous swimming assay (to find symmetric waveform frequency; Fig. 4, Fig. S7, Fig. S11) and microscope chemotaxis assay gradient (to find the relative frequency of symmetric waveform Fig. 6).

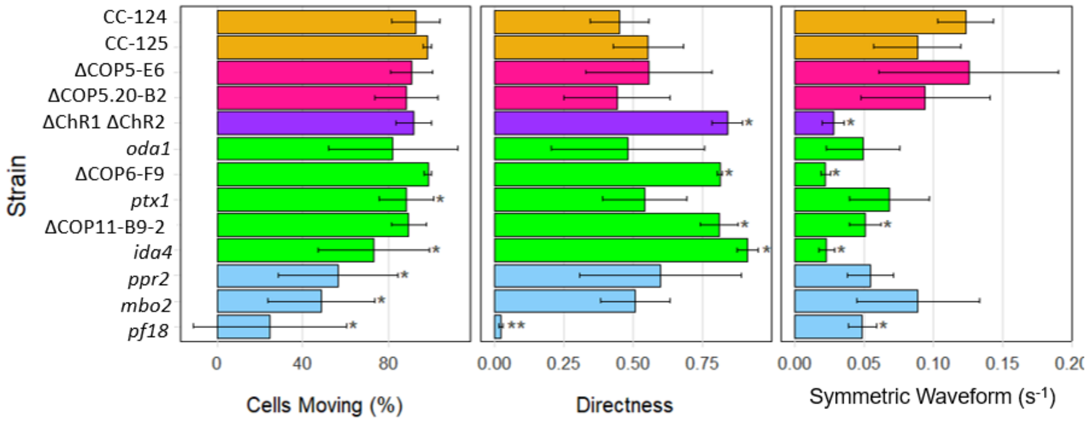


**Fig. S7. Cell moving percentage and directness do not correlate to chemotactic behavior.** A homogenous (non-gradient) swimming assay was performed by suspending cells in TAP and tracking them for 60 seconds. "Cells Moving (%)", shows the percentage of motile cells (defined as having a speed > 15 µm · $sec^{-1}$). "Directness" measures how direct a cell's path is from its initial to final position, calculated by taking the total displacement of a trajectory divided by the overall displacement. "Symmetric Waveform ($s^{-1}$)" represents the frequency of the symmetric waveform of a trajectory per second. Bars colored by the phenotypic cluster as identified in Fig. 3C. Bars represent the average metric from at least 3 trials. Error bars represent standard deviation and P values calculated for each strain compared to CC-125 using a Student's t-test, **p<0.01**,** *p<0.1. Note that the metrics in this figure are calculated from the same experimental dataset used to find the swimming speed of the homogenous swimming assay in Fig. 5.

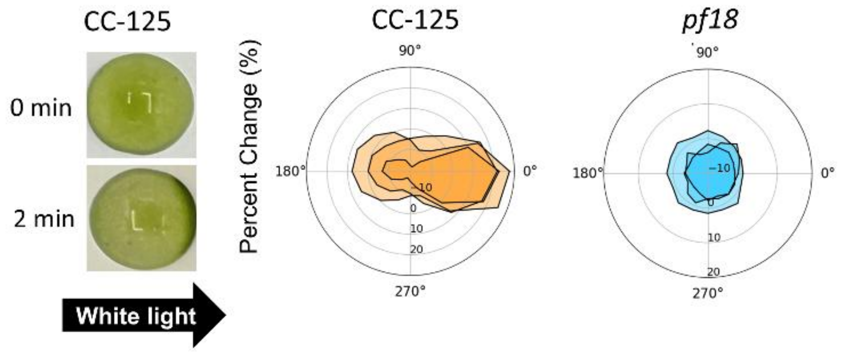


**Fig. S8. Phototaxis droplet assay of a positive (CC-125) and negative control (*pf18*).** Droplet images show one replicate of CC-125 before ($\mathbf{t} = \mathbf{0}$ min) and after ($\mathbf{t} = \mathbf{2}$ min) exposure to a white light gradient from the left-hand side of each droplet. Rose plots show the percent change in image intensity from $\mathbf{t} =$ 2 to $\mathbf{t} =$ 0 for CC-125 and *pf18* of 3 independently performed experiments. The Phototaxis Index (PI) for each experiment is found by taking the standard deviation of the percent change across all quadrants.

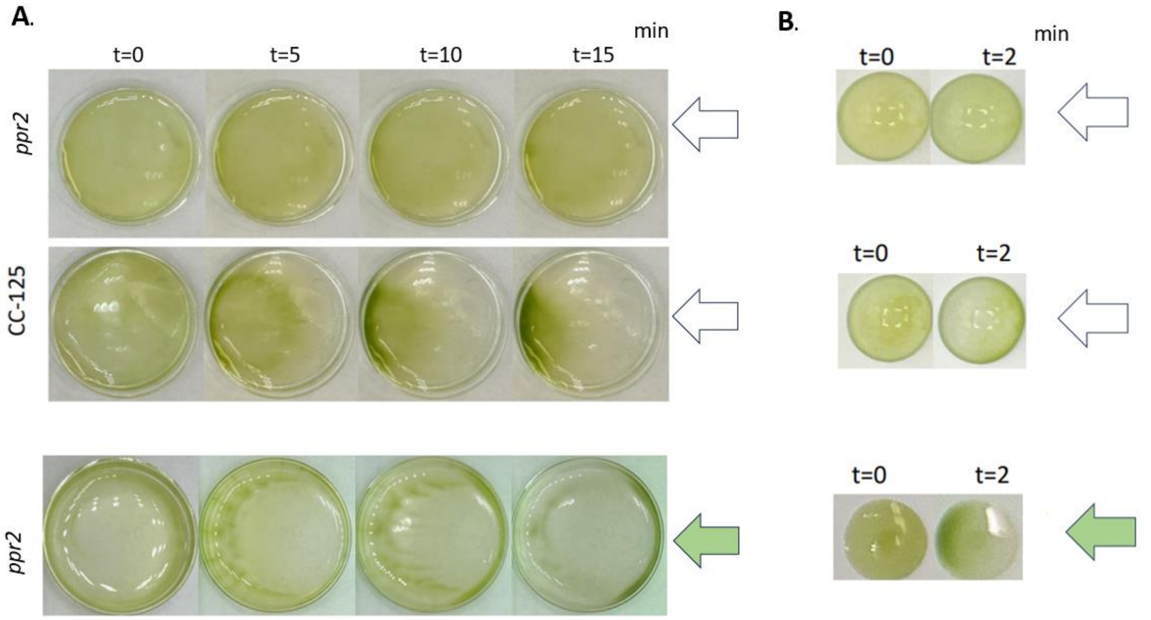


**Fig. S9**. **Conditional effects of light color and assay methods on phototaxis response**. **A. Petri dish phototaxis assay**. The assay was performed in a 3.5 cm diameter Petri dish in the phototaxis assay solution (39) over the course of 15 minutes, illuminated by a white or green light gradient from the right side of the dish (shown as a white and green arrow, respectively). **B. Droplet phototaxis assay**. The assay was performed in 40 μl droplets with the phototaxis assay solution (39) for 2 minutes, illuminated by a white or green light gradient from the right side of the dish (shown as a white and green arrow, respectively). Notice the wild-type strain CC-125 shows a negative phototactic response (the cells migrate against the light source) in the Petri dish assay. In contrast, it shows a positive phototactic response (the cells migrate toward the light source) in the droplet assay. The strain *ppr2* rarely responds to the white light in the Petri dish and droplet assay.

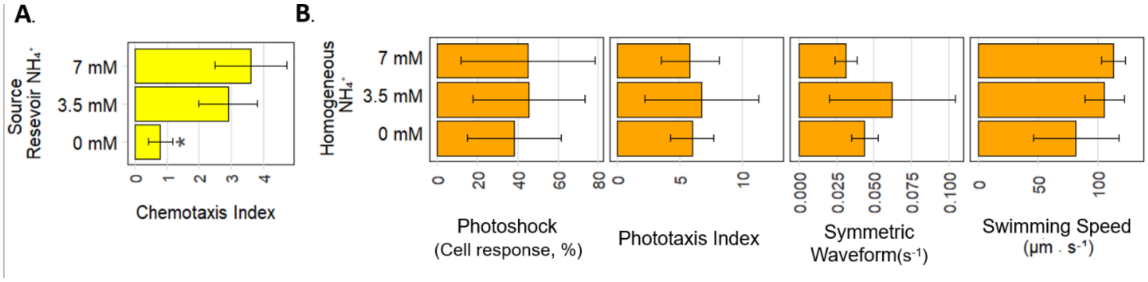


**Fig. S10. *Chlamydomonas* migration toward ammonium source is not conducted by ortho- or klino-kinesis. A. Chemotaxis Lane Assay.** The wild-type strain CC-125 was assayed with different concentrations of $NH_4^+$. Error bars represent standard deviation and P values calculated by comparing to 7 mM $NH_4^+$ using a Student's t-test where (*) $p<0.05$ **B. Analyses of photoshock response, phototaxis, frequency of the symmetric waveform, and swimming speed**. The wild-type strain CC-125 was assayed with different concentrations of $NH_4^+$ in the homogeneous (non-gradient) condition. Error bars represent standard deviation and P values calculated by comparing to 7 mM $NH_4^+$ using a Student's t-test where (*) $p<0.05$. Notice no significant differences were observed among the different homogenous concentrations of ammonium.

**Supplement Methods**

**Bromophenol blue diffusion and gradient modeling.** The source reservoir of the plate is loaded with 200 µm Bromophenol blue (BPB) separated by a 1.5% agarose barrier loaded with 200 µm BPB. The sink reservoir is loaded with deionized water separated by a BPB-free 1.5% agarose hydrogel. 25 inner lanes of the plate filled with 1 ml of deionized water. Timelapse of diffusion of BPB into each lane taken every 30 minutes for 24 hours. Images were imported into ImageJ and converted to 8-bit greyscale, and the vertical intensity profile of each lane at each time point was found (23). Intensity profiles of each lane average together, then normalized to 1, with the highest concentration of BPB being at the source, being equal to 1, and the lowest concentration being at the sink at $t = 0$, to result in normalized intensity (AU). Assuming the agarose reservoir is an infinite source of stable concentration, the experimental data is curve fitted using a custom Python code to a derivative of Ficks Second Law (Equation S1) to find the diffusion coefficient (D) per time point (where $c$ is intensity, $c_0$ is initial intensity, $x$ is position and $t$ is time) as previously described (21, 24, 25). The diffusion coefficient at each time point is then averaged to find the experimental diffusion coefficient of BPB.

$$c(x,t) = c_o * \left(1 - \text{erf}\frac{x}{2\sqrt{Dt}}\right) \quad \text{(Equation S1)}$$

**Chemotaxis index determination.** To calculate the Chemotactic Index (CI) in the Chemotaxis Lane Assay (CLA), a modified method is derived from Sekiguchi et al. (26). Before processing, the outer eighth of each lane is discarded to limit edge aberration effects. Next, the pixel intensity of each lane is divided into twenty equal bins along the 60 mm length of the lane. For each bin, the average pixel intensity at each time point is calculated. To correct for any differences in initial cell concentration per bin, the average pixel intensity of each bin and time point is divided by the value of the corresponding bin at *t = 0*.

The Percent Change from Mean *(PCM)* for each bin and time point was calculated as follows, where $t =$ time, $j =$ bin, and $l =$ time normalized averaged pixel intensity.

$$PCM_{tj} = 100 * \left(1 + \left(1 - \frac{5L_{tj}}{\sum_{j=1}^{5} L_t}\right)\right) \quad \text{(Equation S2)}$$

The Slope Index *(SI)* is found by curve fitting the slope of the *PCM* across all bins for each timepoint to the linear equation.

$$PCM_{tj} = SI_t t * j + b \quad \text{(Equation S3)}$$

The Chemotaxis index (*CI*) was found as the maximum value of the Slope Index across the 24-hour experiment duration.

$$CI = \max(SI_t) \quad \text{(Equation S4)}$$

**Phototaxis droplet assay.** The phototaxis droplet assay was conducted as previously described, with modifications (27). 40 µL of cells suspended in TAP or phototaxis assay solution (28) were pipetted onto a white polypropylene surface. Droplets were placed under homogeneous dim red light (>5 µmol $m^{-2}s^{-1}$) for 10 minutes, then imaged. The droplets were exposed to unidirectional dim white light (>5 µmol $m^{-2}s^{-1}$) for 2 minutes, then imaged. Using ImageJ, each circular droplet was divided into 16 equal radial quadrants (with 1 and 16 being closest to light and with 8 and 9 being farthest). Average pixel intensity per quadrant taken at $t = 0$ and $t = 2$ and used to calculate percent change in pixel intensity after 2 minutes for each quadrant. The standard deviation of the percent change in pixel intensity across all quadrants is calculated, where a larger standard deviation represents stronger phototaxis and a smaller standard deviation represents weaker phototaxis, resulting in the final Phototaxis Index.

**Phototaxis Petri dish assay.** The phototaxis petri dish assay was conducted as previously described, with modifications (27). 2 ml of cells suspended in the phototaxis assay solution (28) were placed in a 3.5 cm diameter Petri dish. The dish was placed under homogeneous dim red light (>5 µmol $m^{-2}s^{-1}$) for 10 minutes and then imaged. The dish was exposed to unidirectional dim white light (>5 µmol $m^{-2}s^{-1}$) or dim green light (>5 µmol $m^{-2}s^{-1}$) for 15 minutes.

**Homogenous (non-gradient) swimming assay.**
A 20 µl *Chlamydomonas* culture was loaded onto a microscope glass with a spacer made from a 3M™ Polyester Film Tape 853 (0.03 mm thickness), and a #1 ½ coverglass was placed on top. To test swimming behavior in non-gradient conditions, cells were suspended in TAP at $1x10^6$ cells · $ml^{-1}$ unless otherwise stated, and timelapse microscopy videos were taken without any light or chemical gradient present during the assay duration. Microscopy was performed using an Olympus IX83 Inverted Microscope equipped with a 20x/N.A.0.8 objective lens and ORCA-Fusion BT CMOS camera. Timelapse images were taken at 30 fps (1497 µm X 1497 µm view field, 2x2 binning) for a 60-second duration. All microscopy was performed in a dark room with a microscope transmission on.

**Photoshock assay.** Cells were incubated in phototaxis assay solution in dim red light for 30 minutes before imaging as described by Okita et. al. (28). During microscopy, the red-light filter (Kodak Wratten filter No. 29) was used to alter the wavelength of the microscope light source, illuminating the cells. Cells were imaged for two seconds before and after white light flash illumination (Canon Speedlite 580EX II, < 1/20 of a second). All other microscope settings were described in the previous section.

**Single-cell tracking and analysis of microscope-based assays.** After video acquisition, cells were tracked with the ImageJ-based plugin TrackMate using the Hessian cell detector and LAP Tracker (29). Tracking yields the *x-y-t* (coordinates and times) for each cell trajectory.

A custom Python code was used to calculate metrics describing each cell trajectory. All trajectories with durations less than 2.5 seconds were removed from the dataset. Cells with an average speed of >10 µm · $s^{-1}$ were classified as non-motile. The number of motile trajectories compared to total trajectories was used to calculate the percent of cells moving. To calculate all other metrics, only motile cell trajectories were used (except for the paralyzed *pf18*, which included non-motile cells).

Five metrics were calculated: cells moving (%), swimming speed (µm · $s^{-1}$), swimming angle (θ), directness, and frequency of the symmetric waveform ($s^{-1}$). In each individual trial, there were numerous cell trajectories, and each individual trajectory had multiple *x-y-t* positional data points (See Movie S3). For directness and symmetric waveform (methodology in Fig. S6), one value was calculated for each individual trajectory. For cell moving, one metric was calculated for the entire cell population in each experiment.

Swimming speed was calculated for every trajectory time point by using the square root of the displacement of the x, and y coordinates in µm at every trajectory time point $t$, compared to the previous frame 0.033 seconds before. The swimming angle was calculated for every trajectory time point based on three x-y-t positions of the cell trajectory (the current timepoint, the timepoint 0.133 secs before, and 0.133 secs after the current timepoint), resulting in angles ranging from 0 to $2\pi$. When specified, the swimming angle is wrapped at 0 to $\pi$, where 0 is a complete reversal and $\pi$ is a straight swimming angle. For metrics calculated at multiple time points and trajectories, the metric is first averaged by trajectory (when applicable) and then averaged by experiment before calculating the reported averages, standard deviations, and statistics.

**Data processing in the microscopic chemotaxis assay.** Single-cell tracking was performed with TrackMate as described above. All tracks shorter than 4 seconds are discarded. First, whether a cell was either in 1) asymmetric or 2) symmetric waveform was determined at 0.033-second intervals according to an algorithm (Fig. S6). Then, the orientation angle of the tracked cells was determined. The orientation angle was defined as the angle (0 to 2π) relative to the source (7 mM $NH_4^+$ agarose block). The x-y coordinate plane was set so that the source and sink were located on the left-hand and right-hand side in the x-axis, respectively. An x-y coordinate of a cell at a given time point was the origin (0,0). An x-y coordinate of the same cell was determined 0.33 seconds after moving from the origin. The angle formed between the origin, the previous coordinate, and the new coordinate was the orientation angle. Hence, in this analysis, an orientation angle of π means a cell moves straight toward the source in 0.33 seconds. An orientation angle of 0 means a cell moves straight toward the sink in 0.33 seconds. The number of asymmetric and symmetric waveform occurred in the 10-second observation period was determined. Due to the video analysis starting at t=60 secs after initial exposure to the ammonium gradient, the wild-type strain CC-125 had more total cell trajectories than the mutant strain ΔCOP5, as cells had begun to accumulate at higher densities near the ammonium source. To correct for the difference in the number of cell trajectories in the two strains, the relative frequency of symmetric waveform to asymmetric waveform in two opposite orientation angles (toward the source π ± π/5 and toward the sink 0 ± π/5) was normalized by the total number of symmetric waveform and asymmetric waveform events as determined for all orientation angles, respectively.

**Movie S1 (separate file). Identification symmetric waveform for individual trajectories in CC-125 photoshock assay.** Movie taken over the course of 4 seconds. Light flash shock at t = 2 sec. Video on right shows cells trajectories found after single cell tracking and processing with Python. Trajectories colored by “symmetric” (green) and “asymmetric” (red), as determined by a custom Python code.  Video on left shows the corresponding microscopy video taken for a 4 second duration at 30 FPS, using 20x magnification. Red light filter used for the duration of microscopy recording.

**Movie S2 (separate file). Cell migration of WT CC-125 in microscopic chemotaxis assay.** Cell swimming of CC-125 imaged with a 30-fps rate (every 0.033 sec) in a flow-free ammonium gradient. Ammonium source to the left-hand side of the movie. Movie captured 60 second after cell transfer the cover glass chamber for a 10 second duration. Movie taken adjacent to the $NH_4^+$ source, (see Fig. 6A schematic), using ring light to provide homogenous white light exposure.

**Movie S3 (separate file). Single Cell tracking of CC-125 in homogeneous swimming assay.** Leftmost movie Shows CC-125 swimming in homogenous TAP medium, taken using 20x/N.A.0.8 objective lens for a 60 second duration, 30-fps rate (every 0.033 sec). The right two graphs show single cell tracking of individual cells over time with initial trajectory position adjusted to 0,0 on the XY coordinate plane. The trajectories of the middle graph are colored by swimming speed and the trajectories of the rightmost graph are colored by swimming angle, according to their respective color lookup tables.

**Movie S4 (separate file). Ammonium chemotaxis and gradient formation in a chambered coverglass.** Images were taken for a duration of two hours, at 5-minute intervals. For the chamber to the left, the chemical source (TAP+N), is the bottom agarose block and the chemical sink is the top agarose block (TAP-N). 1 ml of algae suspended in TAP-N was incubated in homogenous white light for a 10-minute duration prior to transferring to the cover glass chamber on the left. The cover glass chamber on the right was filled with deionized water, to show gradient formation over time. Scale bar shown on movie.
