## Supplementary figures and images for "Membrane-bound Guanylyl Cyclase COP5/HKR1 changes ciliary beat pattern and biases cell steering during chemotaxis in *Chlamydomonas reinhardtii*"

### Supplement Movie 1

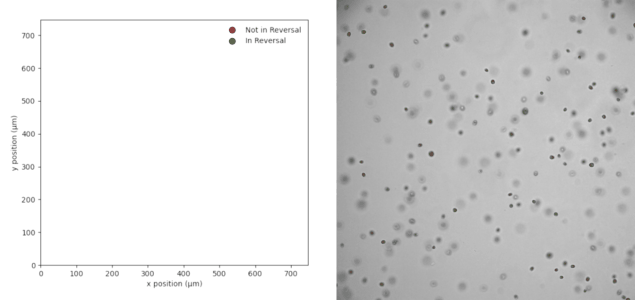
